# *Ade2* functions in the *Drosophila* fat body to promote sleep

**DOI:** 10.1101/361055

**Authors:** Maria E. Yurgel, Kreesha D. Shah, Elizabeth B. Brown, Ryan A. Bennick, Justin R. DiAngelo, Alex C. Keene

## Abstract

Metabolic state is a potent modulator of sleep and circadian behavior and animals acutely modulate their sleep in accordance with internal energy stores and food availability. Across phyla, hormones secreted from adipose tissue act in the brain to control neural physiology and behavior to modulate sleep and metabolic state. Growing evidence suggests the fat body is a critical regulator of complex behaviors, but little is known about the genes that function within the fat body to regulate sleep. To identify molecular factors functioning in the periphery to regulate sleep, we performed an RNAi screen selectively knocking down genes in the fat body. We found that knockdown of *Phosphoribosylformylglycinamidine synthase*/*Pfas* (*Ade2*), a highly conserved gene involved the biosynthesis of purines, reduces sleep and energy stores. Flies heterozygous for multiple *Ade2* mutations are also short sleepers and this effect is partially rescued by restoring *Ade2* to the fat body. Targeted knockdown of *Ade2* in the fat body does not alter arousal threshold or the homeostatic response to sleep deprivation, suggesting a specific role in modulating baseline sleep duration. Together, these findings suggest *Ade2* functions within the fat body to promote both sleep and energy storage, providing a functional link between these processes.

## Introduction

Animals balance nutritional state and energy expenditure in order to achieve metabolic homeostasis (Taguchi and White 2008; Yurgel *et al*. 2015). In the fruit fly, *Drosophila melanogaster*, feeding behavior and metabolism are regulated by peripheral tissues including muscle, liver, the adipose-like organ called the fat body, and the gastrointestinal tract (Erion and Sehgal 2013; Itskov and Ribeiro 2013). Similarly, in mammals, endocrine hormones such as ghrelin, leptin, insulin, and glucagon are secreted from the stomach, adipose, and pancreas, to convey nutritional status to the brain regions that regulate sleep and metabolism (Marks *et al*. 2009; Bruijnzeel *et al*. 2011; Karra *et al*. 2013) Dysregulation of peripheral hormonal signals leads to a number of metabolic diseases including obesity, diabetes, and insomnia (Arble *et al*. 2015). Therefore, mechanistic investigation of factors regulating brain-periphery communication is critical to understanding disorders associated with sleep and metabolism.

Adipose tissue senses overall nutrient levels in the animal and modulates hunger-induced behaviors through controlling energy storage and secreting factors that act on the nervous system to affect behavior (Xu *et al*. 2011a; Sassu *et al*. 2012; Musselman *et al*. 2013). The insect fat body is central to the control of energy homeostasis. It is the primary site of glycogen and triglyceride storage, and is the main detoxification organ in the fly, thereby exhibiting functions analogous to the mammalian liver and adipose tissue (Arrese and Soulages 2010). Genome-wide transcriptome analysis identified many genes upregulated during starvation, including metabolic enzymes, cytochromes, metabolite transporters, kinases, and proteins involved in lipid metabolism (Grönke *et al*. 2005). Even though much is known about the primary function of these genes in regulating energy storage, little is known about how they may impact sleep and other behaviors.

Many of the genes and transmitters required for metabolic regulation of sleep and feeding in mammals are conserved in *Drosophila* (Allada and Siegel 2008; Padmanabha and Baker 2014), and numerous conserved factors have been identified that regulate sleep and metabolic function (Arble *et al*. 2015; Yurgel *et al*. 2015). The GAL4/UAS system, in combination with genome-wide RNAi libraries, allow for selectively decreasing gene expression in the fly fat body and then measuring the effects on sleep (Brand and Perrimon 1993; Dietzl *et al*. 2007). Growing evidence suggests the fat body regulates complex behaviors including sleep (Lazareva *et al*. 2007; Kim *et al*. 2017; Umezaki *et al*. 2018), yet the molecular basis through which the fat body regulates sleep remains poorly understood. Here, we sought to identify sleep regulators in the fat body by selectively decreasing the expression of genes that have been previously identified to be upregulated in the fat body of starved flies, and measuring their effects on sleep (Grönke *et al*. 2005).

We identified *Phosphoribosylformylglycinamidine synthase (Ade2)*, a highly conserved gene involved the biosynthesis of purines, as required for normal sleep in flies. Flies deficient for *Ade2* are short-sleepers and have reduced triglyceride stores, suggesting that loss of *Ade2* impairs energy storage and inhibits sleep. Disruption of *Ade2* in the fat body does not disrupt arousal threshold and homeostatic response to sleep deprivation, suggesting a specific role in modulating baseline sleep duration. These findings provide a novel factor that functions in the fat body to regulate sleep, and support growing evidence that peripheral metabolic tissue is critical for the proper regulation of sleep.

## Results

To identify genes expressed in adipose tissue that regulate sleep, we performed an RNAi screen by assaying the TRiP RNAi collection to selectively knock down genes in the fat body (Brand and Perrimon 1993; Ni *et al*. 2009). A total of 113 genes previously reported to be upregulated in whole flies during starvation (Grönke *et al*. 2005) were selectively knocked down in the fat body using the GAL4 driver, CG-GAL4 (Asha *et al*. 2003), and female flies were then assayed for sleep (Fig 1A) in *Drosophila* Activity Monitors (Pfeiffenberger *et al*. 2010a). Flies with RNAi targeted to the fat body were compared to controls expressing RNAi targeted to luciferase (CG-GAL4>luc-RNAi). Knockdown of *Ade2* in the fat body (CG-GAL4>*Ade2*-RNAi) resulted in a loss of over 200 minutes of sleep, while a knockdown of the glucose transporter *CG6484* (CG-GAL4>*CG6484*-RNAi) and *CG6767*, a kinase involved in purine/pyrimidine metabolism, (CG-GAL4> *CG6767*-RNAi) resulted in increased sleep (Table 1). We chose to focus analysis on the role of *Ade*2 in sleep regulation because of the robust sleep loss phenotype identified in the screen. To verify these results, we retested the effects of *Ade2* knockdown on sleep. In agreement with the screen, sleep was reduced in *Ade2* knockdown flies (CG-GAL4> Ade2-RNAi) compared to control flies with CG-GAL4 driving RNAi targeted to luciferase (CG-GAL4>luc-RNAi) (Fig 1B). Quantification of sleep throughout the 24 hr testing period revealed that fat body specific knockdown of *Ade2* results in sleep loss with significant reductions during both the day and night periods, suggesting *Ade2* is required for both day and nighttime sleep (Fig 1B,C).

**Fig 1.**
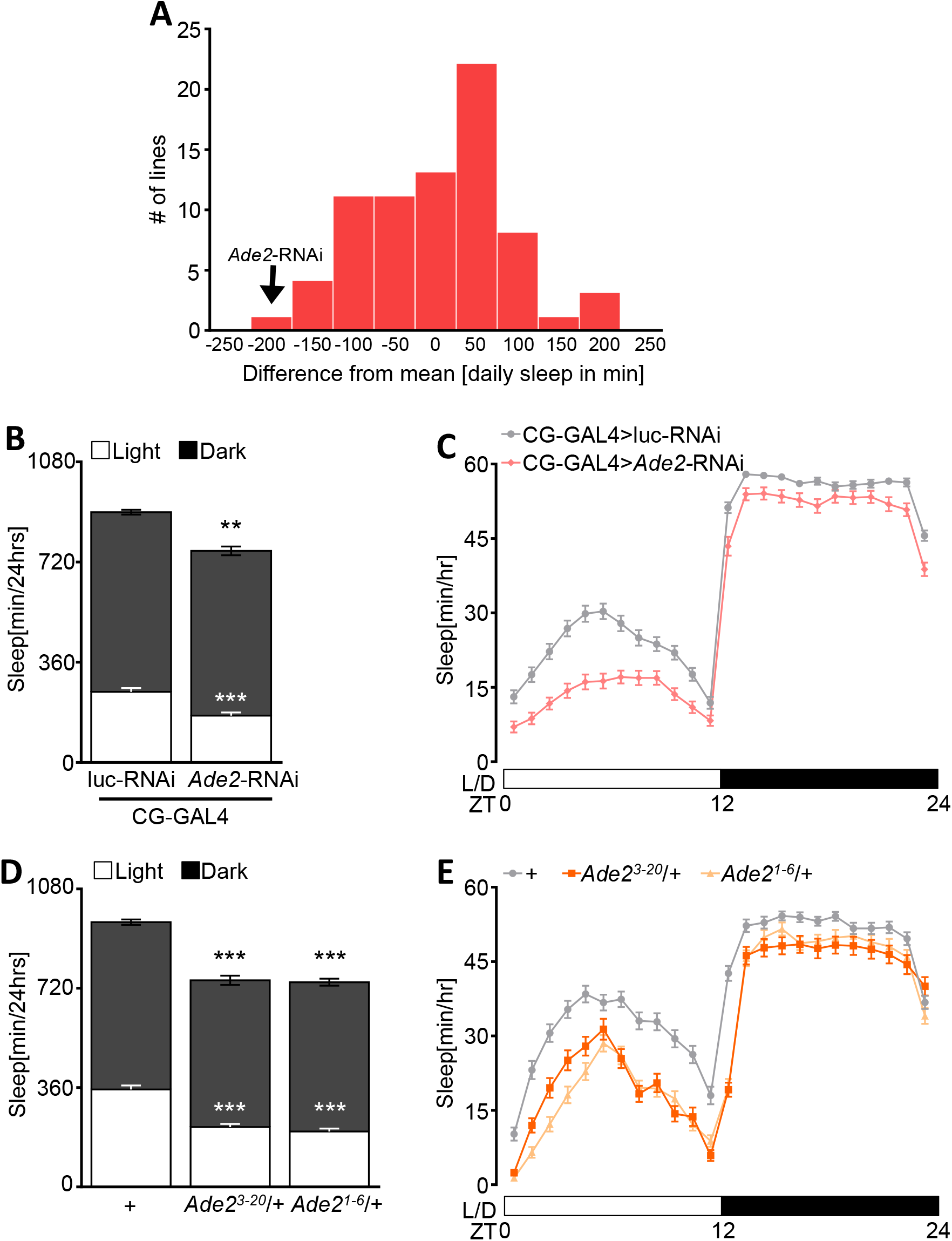
Ade2 functions in the fat body to promote sleep. (**A**) Histogram showing the distribution of sleep over 24 hours from fat body-specific knockdown of genes previously reported to be upregulated during starvation. Daily sleep is depicted as the difference between the mean of a group of ~80 viable lines tested. Black arrow indicates the *Ade2-*RNAi control line. (**B**) Knock down of *Ade2* in the fat body (CG-GAL4>UAS-*Ade2-*RNAi; n=84) results in a significant reduction in sleep during daytime (white; *p*<0.0001, t=6.241) and nighttime (black; *p*=0.0004, t=3.614) compared to control flies (CG-GAL4>UAS-luc-RNAi; n=115). Unpaired t-test. (**C**) Sleep profile of hourly sleep averages over a 24 hour experiment. White/black bars represent lights on and off, respectively. ZT denotes Zeitgeber time. Sleep is reduced in flies expressing *Ade2*-RNAi in the fat body (pink) compared to control (grey). (**D**) Sleep is significantly reduced in *Ade2^3-20^*/+ mutants (n=66) and *Ade2^1-6^*/+ (n=90) during daytime (*p*<0.0001 for all groups) and nighttime (*p*=0.0002 *and p*=0.004) compared to *w^1118^* control (n=110). One-way ANOVA, Light, F(2, 261)=11.20; Dark, F(2, 263)=45.91. (**E**) Sleep profile of hourly sleep for *Ade2^3-20^/+* mutants (dark orange), *Ade2^1-6^/+* (light orange), and *w^1118^* control (grey). All columns are mean ± SEM; \**p*<0.05; \*\**p*<0.01; \*\*\**p*<0.001.

To confirm that the sleep loss phenotype observed with *Ade2* knockdown was not due to off-target effects of RNAi, we assayed sleep in *Ade2* mutant *Drosophila*. Two *Ade2* mutants, *Ade2^3^*-^20^ and *Ade2^1-6^*, have been generated by *P*-element excision (Holland *et al*. 2011). For *Ade2^1-6^*, the deletion includes the transcription start site and part of the first coding exon resulting in a null allele (Holland *et al*. 2011). While both alleles are homozygous lethal, heterozygous flies are viable, flies heterozygous for *Ade2*^3-20^ or *Ade2*^1-6^ sleep less than *w^1118^* flies (the background control strain) during the day and night, phenocopying results obtained with RNAi (Fig 1D). Further, a sleep profile analysis confirms sleep loss is reduced throughout the day and night confirming the sleep phenotype observed in RNAi knockdown flies (Fig 1E).

Reduced sleep can be accounted for by a reduction in the total number of sleep bouts, shortened duration of individual sleep bouts, or a combination of both (Garbe *et al*. 2015a). RNAi knockdown of *Ade2* (CG-GAL4>UAS-*Ade2*-RNAi) resulted in reduced sleep bout length compared to control flies (Fig S1A), while total sleep bout number was reduced during the day and increased during the night (Fig S1B). Similarly, average sleep bout length was significantly reduced in both *Ade2* mutants (*Ade2^3-20^* and *Ade2^1-6^*) compared to controls, suggesting that both *Ade2*-RNAi and mutant flies present a less consolidated sleep pattern (Fig S1C). No difference in sleep bout number was detected during the daytime in flies heterozygous for the *Ade2^3-20^* or *Ade2^1-6^* mutations compared to *w^1118^* controls, while a significant increase in sleep bout number was detected for both heterozygous mutants during the night (Fig S1D). Taken together, these experiments suggest that the short sleeping phenotype of *Ade2* deficient flies primarily derives from the shortening of sleep bouts.

To determine whether the sleep phenotype might be explained by generalized changes in locomotor activity, we analyzed waking activity (beam breaks/minute) in *Ade2* knockdown flies and flies heterozygous for each mutation. Waking activity did not significantly differ between CG-GAL4>*Ade2*-RNAi flies compared to control flies, though CG-GAL4>*Ade2*-RNAi trended toward increased waking activity during the day and the night (Fig S1E). Similarly, waking activity was significantly increased in *Ade2*^1-6^ heterozygous flies during daytime and nighttime compared to *w^1118^* controls, while *Ade2*^3-20^/+ had a significant increase in daytime but not nighttime waking activity (Fig S1F). Together, these findings suggest that disruption of *Ade2* function induces hyperactivity, in addition to shortening sleep phenocopying starved flies.

To verify that expression of *Ade2* in the fat body was sufficient for normal sleep, we selectively restored *Ade2* to the fat body in the background of *Ade2* heterozygous flies and measured sleep. Fat body rescue flies (CG-GAL4>UAS-*Ade2*; *Ade2*^3-20^/+) slept more than *Ade2^3-20^*/+ heterozygous mutants harboring the UAS-*Ade2* transgene without the GAL4 (UAS-*Ade2*; *Ade2^3-^*^*20*^/+) (Fig 2A,B). However, fat body expression did not fully rescue sleep, as rescue flies slept less than control flies harboring CG-GAL4 or UAS-*Ade2* transgenes alone. Therefore, restoration of *Ade2* to the fat body partially restores sleep to *Ade2*^3-20^ mutant flies. Similarly, restoring *Ade2* within the fat body of flies heterozygous for the *Ade2*^1-6^ mutation (CGGAL4>UAS-*Ade2*; *Ade2*^1-6^/+) partially restores sleep, with rescue flies sleeping significantly more than UAS-*Ade2*; *Ade2*^3-20^/+ heterozygous flies, but less than flies harboring CG-GAL4 transgene alone (Fig 2C,D). Expression of *Ade2* in the fat body of flies heterozygous for *Ade2*^3-20^ or *Ade*2^1-6^ rescued both average sleep bout length during nighttime and sleep bout number during the day and nighttime to control levels (Fig S2A-D). Similarly, the hyperactivity phenotype seen in *Ade*2^3-20^ and *Ade2*^1-6^ heterozygous mutant flies was restored in rescue flies, and these flies did not differ from heterozygous controls (Fig S2E, S2F). To determine whether upregulation of *Ade2* in the fat body is sufficient to promote sleep, we overexpressed *Ade2* in the fat body of wildtype flies (CG-GAL4>UAS-*Ade2*). Sleep in these flies did not differ from transgenic controls harboring CG-GAL4 or UAS-*Ade2* alone (Fig 2E). Taken together, these findings suggest *Ade2* expression in the fat body is necessary for normal sleep, but enhanced *Ade2* expression is not sufficient to increase sleep.

**Fig 2.**
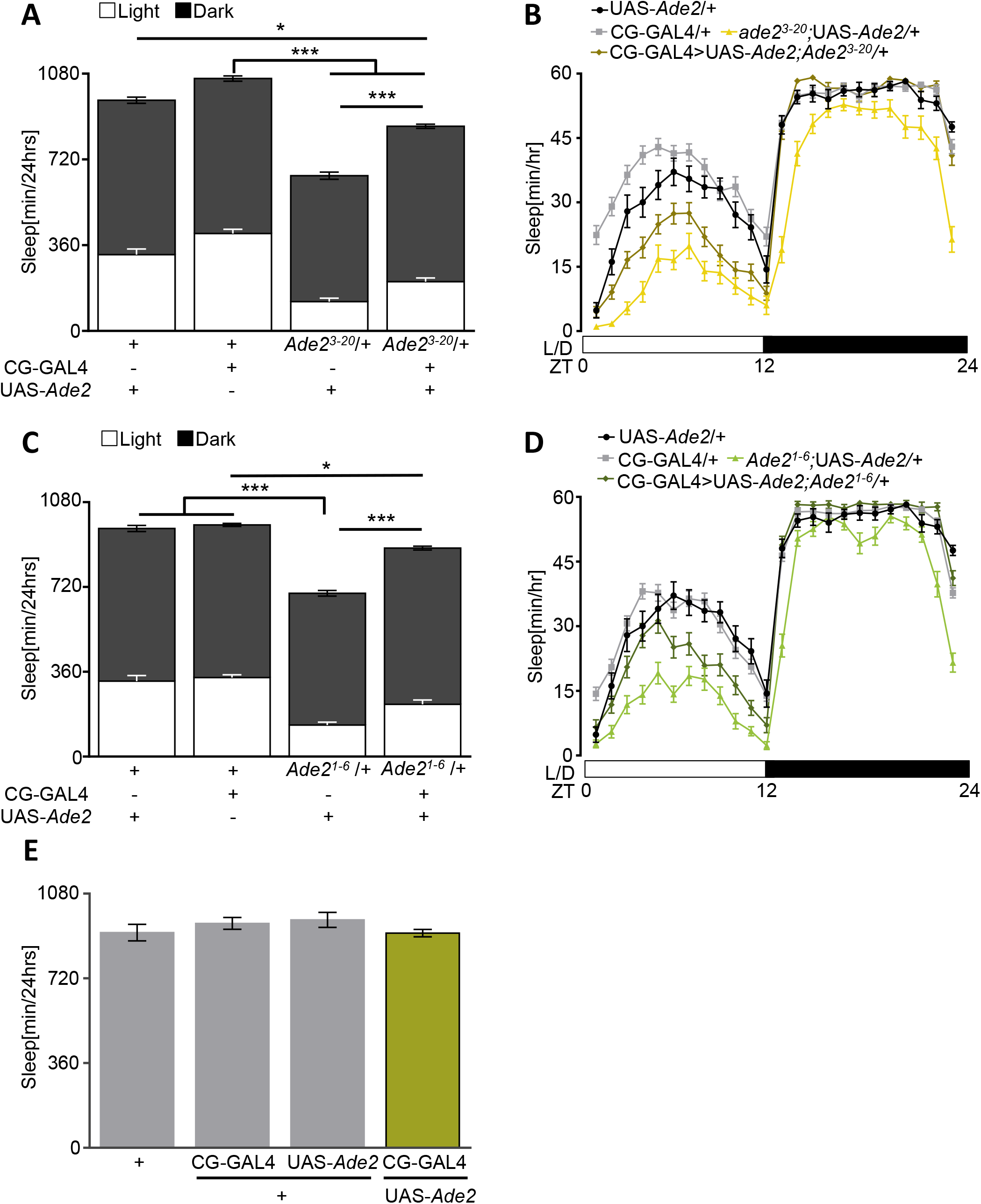
*Ade2* expression in the fat body partially rescues sleep loss. (**A**) Fat body rescue of *Ade2^3-20^/+* (CG-GAL4>UAS-*Ade2;Ade2^3-20^*/+; n=43) partially restores total sleep compared to CG-GAL4/+ (n=69, *p*<0.0001) control and UAS-*Ade2*/+ (n=31, *p*=0.014). Total sleep duration in rescue flies is significantly increased compared to *Ade2^3-20^*/+;UAS-*Ade2* mutant control (n=30, *p*<0.0001). One-way ANOVA, F(3, 169)= 37.76. (**B**) Sleep profile of hourly sleep averages over a 24 hour experiment for *Ade2^3-20^* rescue (gold) compared with *Ade2^3-20^*/+;UAS-*Ade2* (yellow), CG-GAL4/+ (grey), and UAS-*Ade2*/+ (black) controls. White/black bars represent lights on and off. ZT denotes Zeitgeber time. (**C**) Total sleep is significantly increased in flies expressing *Ade2* in the fat body of *Ade2^1-6^* mutants (n=37) compared to *Ade2^1-6^*/+;UAS-*Ade2* mutant controls (n=39, *p*<0.0001). Rescue flies are significantly different than control CG-GAL4/+ (grey, n=79, *p*=0.010). *Ade2^1-6^*/+;UAS-*Ade2* mutants have reduced sleep compared to UAS-*Ade2*/+ (n=31, *p*<0.0001) and CG-GAL4/+ (*p*<0.0001) controls. One-way ANOVA, F(3, 182)= 41.94. (**D**) Sleep profile of hourly sleep averages over a 24 hour experiment for *Ade2^1-6^* rescue (dark green) compared with *Ade2^1-6^*/+;UAS-*Ade2* (light green), CG-GAL4/+ (grey), and UAS-*Ade2*/+ (black) controls. (**E**) Total sleep did not differ between flies overexpressing *Ade2* in the fat body (CG-GAL4>UAS-*Ade2;* n=64) compared to *w^1118^* (n=55, p>0.99), CG-GAL4/+(n=32, p=0.74), or UAS-*Ade2*/+ (n=31, p=0.51) controls. One-way ANOVA, F(3,178)=0.43. All columns are mean ± SEM; \**p*<0.05; \*\**p*<0.01; \*\*\**p*<0.001.

Sleep is associated with an elevated arousal threshold where animals are less responsive to environmental stimuli (Campbell and Tobler 1984; Hendricks *et al*. 2000; Shaw *et al*. 2000). To determine whether *Ade2* regulates arousal threshold, we measured sleep in the ***D**rosophila* **AR**ousal **T**racking system (DART, Fig 3A) (van Alphen *et al*. 2013; Faville *et al*. 2015). Briefly, the system allows for automated video-tracking combined with controlled application of a vibration stimulus. The response of sleeping animals to the vibration is used to determine the arousal threshold (Fig 3A). In agreement with infrared-based recordings, video-monitoring in the DART system confirmed reduced sleep in CG-GAL4>*Ade2*-RNAi flies with *Ade2* knocked down in the fat body (Fig 3B,C). No differences in arousal threshold were detected between *Ade2* knockdown and control flies during the daytime or nighttime suggesting *Ade2* affects sleep duration, but not sleep-associated changes in arousal (Fig 3D). Similarly, video monitoring in the DART system confirmed sleep duration was reduced in *Ade2*^3-20^ and *Ade2*^1-6^ heterozygous flies, but no effect on arousal threshold was detected during the day or night (Fig 3E-G). Together, these results suggest that that arousal threshold is not altered in *Ade2* deficient flies.

**Fig 3.**
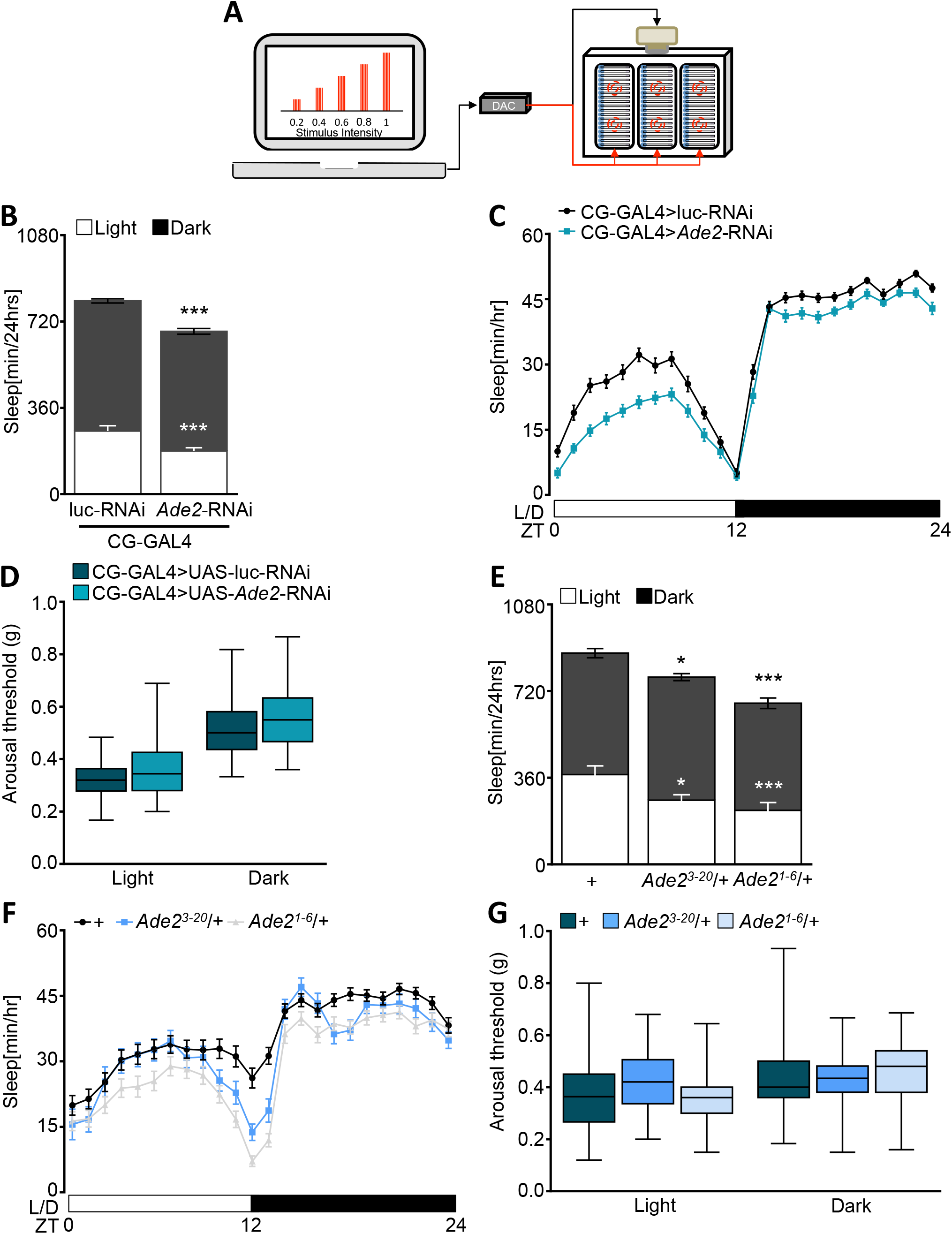
Arousal threshold is normal in *Ade2*-RNAi and *Ade2* mutants. (**A**) The *Drosophila* Arousal Tracking (DART) software records fly movement while simultaneously controlling mechanical stimuli via a digital analog converter (DAC). Mechanical stimuli are delivered to three platforms, each housing twenty flies under the control of two motors. Mechanical stimuli of increased strength were used to assess arousal threshold (shown on the computer screen). Arousal thresholds were determined hourly, starting at ZT = 0. (**B**) Video-tracking analysis of sleep. Sleep during daytime (white, *p*<0.0001, t=6.11) and nighttime (black, *p*<0.0001, t=5.47) is significantly reduced in flies expressing *Ade2*-RNAi in the fat body (CG-GAL4>UAS-*Ade2*-RNAi; n=105) compared to control (CG-GAL4>UAS-luc-RNAi; n=114). Unpaired t-test. (**C**) Sleep profile over a 24 period. White/black bars represent lights on and off. ZT denotes Zeitgeber time. *Ade2* knock down flies (turquoise) sleep less than control (black). (**D**) Arousal threshold during dayttime (*p*=0.06) and nighttime (*p*=0.07) does not differ between CG-GAL4>UAS-*Ade2*-RNAi (turquoise; n=79) and control (dark green;n=85). Mann-Whitney U; Light, 2783; Dark, 2807. (**E**) Video-tracking analysis shows reduced sleep during daytime (*p*=0.045) and nighttime (*p*=0.034) in *Ade2^3-20^*/+ (n=46) compared to *w^111^* control flies (n=87). *Ade2^1-6^*/+ mutant flies (n=80, *p*<0.0001) sleep significantly less than control flies during day and night (*p*<00001). One way-ANOVA, Light, F(2,210)=15.11; Dark, F(2,210)=17.94 (**F**) Sleep profile representative of data in (E). *w^1118^* control flies (black) slept more than *Ade2^3-20^*/+ (blue) and *Ade2^1-6^*/+ (grey) mutants. (**G**) Arousal threshold does not differ between *w^1118^* control (dark green n=71) and *Ade2^3-20^*/+ (blue, n=29) and *Ade2^1-6^*/+ (light blue; n= 63) during light and dark. Kruskall Wallis; Light, 6.87; Dark, 3.02. All columns are mean ± SEM; \**p*<0.05; \*\**p*<0.01; **-\**p*<0.001.

Growing evidence suggests that independent neural mechanisms regulate sleep under undisturbed conditions and the homeostatic sleep rebound following deprivation (Seidner *et al*. 2015; Liu *et al*. 2016). To determine if sleep homeostasis is intact in *Ade2* deficient flies, we sleep deprived flies by mechanical shaking for 12-hours throughout the night, and measured sleep during the following day. The sleep deprivation protocol resulted in in significant sleep rebound in flies heterozygous for the *Ade2*^3-20^ and *Ade2*^1-6^ mutations, similar to controls (Fig 4A,B). Together, these results suggest *Ade2* is dispensable for homeostatic sleep rebound.

**Fig 4.**
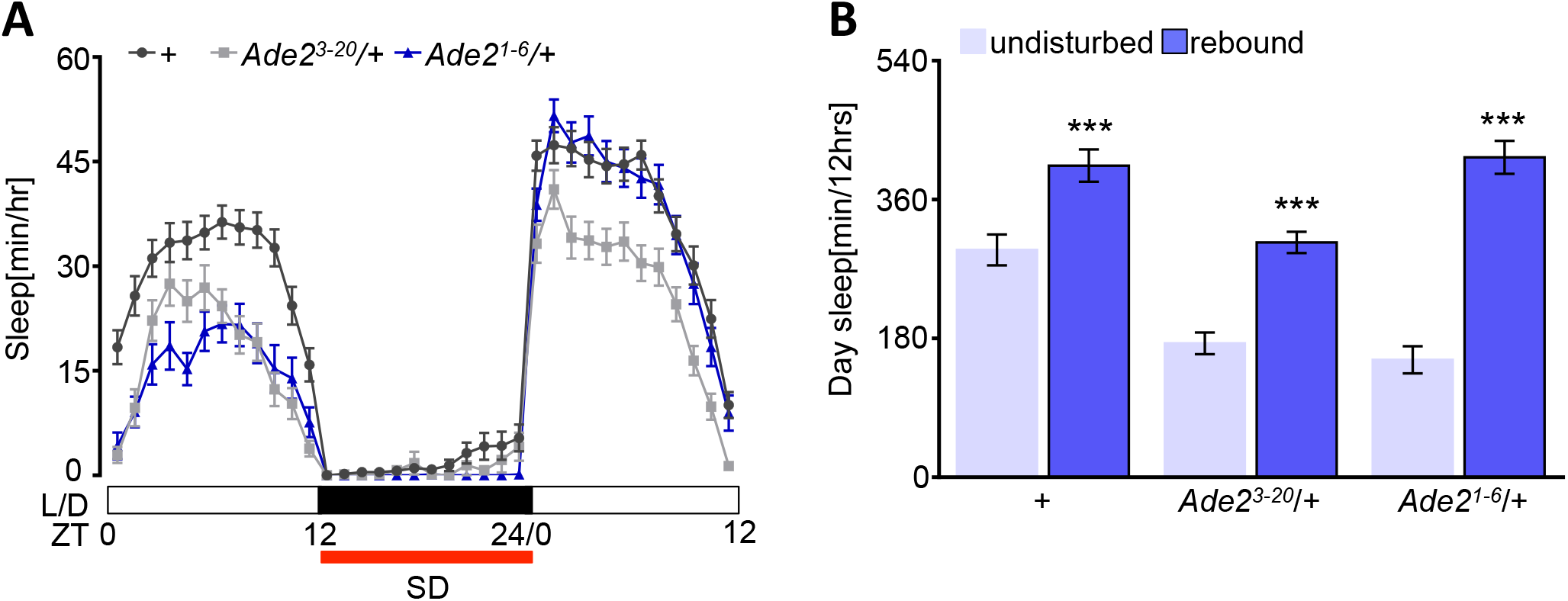
Homeostatic recovery sleep is not altered in *Ade2* mutants. (**A**) Sleep profile for hourly sleep for 36 hours. Flies are undisturbed on the first 12 hours (ZT0-ZT12), lights on (white bars). In the subsequent night (black bars, ZT12-ZT24), flies are mechanically sleep deprived (red), and rebound is measured in the next 12 hours during the light period (ZT0-ZT12). ZT denotes Zeitgeber time. (**B**) *w^1118^* control (n=78, *p*=0.0001), *Ade2^3-20^*/+ (n=51, *p*=0.0002), and *Ade2^1-6^*/+ mutants (n=50, *p*<0.0001) significantly rebound after sleep deprivation (purple) compared to undisturbed day (light purple). Two-way ANOVA, F(2, 352)=8.75. All columns are mean ± SEM; \**p*<0.05; \*\**p*<0.01; \*\*\**p*<0.001.

Flies suppress sleep and increase waking activity in response to starvation and when energy stores are depleted; therefore, we reasoned that the loss of sleep in *Ade2* mutants may be due to reduced energy storage (Lee and Park 2004; Keene *et al*. 2010; Murakami *et al*. 2016). To determine if energy stores are dysregulated in *Ade2* mutants, we measured whole-body triglycerides, glycogen, and free glucose levels in fed *Ade2* loss of function flies using colorimetric assays (Birnbaum *et al*. 2012). Knockdown of *Ade2* selectively in the fat body (CGGAL4>*Ade2*-RNAi) resulted in reduced triglycerides and free glucose levels (Fig 5A-C) without affecting glycogen levels. Similarly, triglyceride and free glucose levels were reduced in *Ade2*^3-20^ heterozygote flies, while glycogen levels were unaffected (Fig 5D-F). In *Ade2*^1-6^ heterozygous flies, only free glucose levels are reduced (Fig 5E). Taken together, these findings suggest *Ade2* function in the fat body is required for the normal storage of triglycerides and free glucose, supporting the notion that the reduced sleep may be caused metabolic changes that place the fly in a starvation-like state.

**Fig 5.**
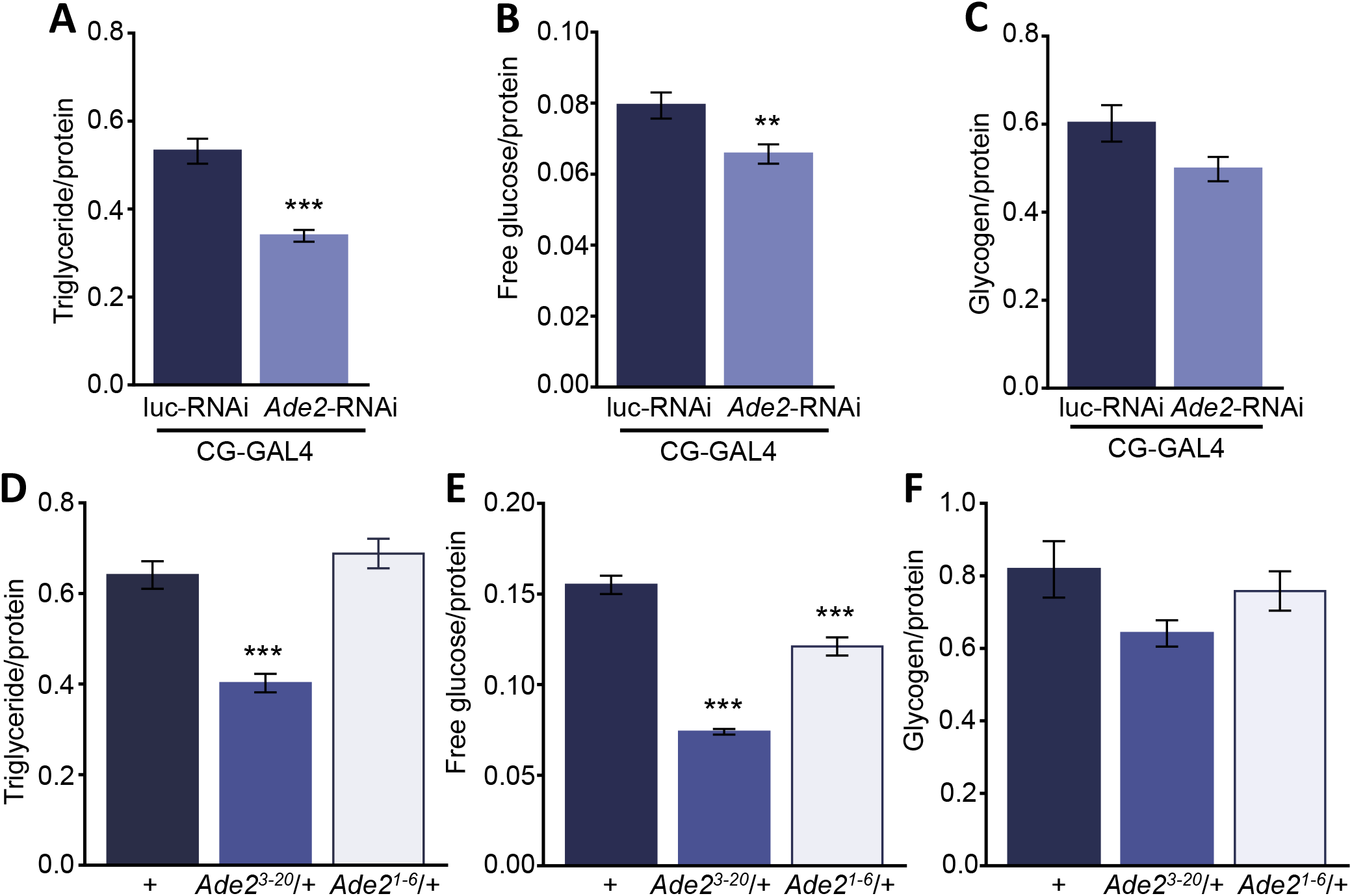
Triglycerides and free glucose are altered in *Ade2* knock down and mutants. (**A-C**) Triglyceride (**A**, *p*<0.001, t=0.56) and free glucose levels (**B**, *p*=0.007, t=0.25) are reduced in flies with *A*de2 knock down in the fat body (CG-GAL4>UAS-*Ade2*-RNAi; n=13) compared to control flies (CG-GAL4>UAS-luc-RNAi; n=15), while there are no significant differences in glycogen levels (**C**, n=13, *p*=0.053, t=2.025). Unpaired t-test. (**D-F**) Triglyceride (**D**, n=14 *p*<0.0001) and free glucose levels (**E**, *p*<0.0001) are reduced in *Ade2^3-20^*/+ mutants (n=14) compared to *w^1118^* controls (n=14), while there are no significant differences in glycogen levels (**F**, *p*=0.75). *Ade2^1-6^* mutants (n=15) show a reduction in free glucose levels (**E**, *p*<0.0001) compared to *w^1118^* controls. Triglyceride (**D**, *p*=0.47) and glycogen (**F**, *p*=0.09) stores do not differ between *Ade2^1-6^* and control flies. One-way ANOVA; TAG, F(2,40)=28.59; Free Glucose, F(2,40)=90.30; Glycogen, F(2,40)=2.45. All columns are mean ± SEM; \**p*<0.05; \*\**p*<0.01; **-\**p*<0.001.

## Discussion

To our knowledge, this study represents the first genetic screen for *Drosophila* sleep regulators that specifically examine the role of non-neuronal tissue in sleep regulation. The fat body is critical for regulating energy storage in *Drosophila* and has been implicated in many behaviors including sleep regulation, courtship, circadian rhythms, and feeding (Lazareva *et al*. 2007; Xu *et al*. 2011; Thimgan *et al*. 2012; Kim *et al*. 2017; Thimgan *et al*. 2010). While the genetic examination of many behaviors, including sleep, have predominantly focused on investigating the neural regulation of behavior, the contribution of the fat body to behavioral regulation is less understood (Iijima *et al*. 2009; Xu *et al*. 2011b; Sassu *et al*. 2012). A complete understanding of behavior will require systematic investigation of the role of the fat body and other peripheral tissues in behavioral regulation.

The fat body-specific screen for sleep regulators described here identified *Ade2* function within this tissue type as critical for normal sleep. These findings add to a growing body of literature suggesting that the adipose tissue is a critical regulator of sleep in both mammals and invertebrates (Laposky 2005; Thimgan *et al*. 2010; Arble *et al*. 2015). In inbred fly lines derived from wild caught *Drosophila*, sleep duration positively associated with whole body triglyceride levels, and *Drosophila* selected for starvation resistance have elevated fat body stores and prolonged sleep (Harbison *et al*. 2004; Masek *et al*. 2014a; Slocumb *et al*. 2015). Similarly, flies mutant for the triglyceride lipase gene *brummer* have elevated triglyceride stores and an enhanced homeostatic sleep response, while mutants for the perilipin-like protein *lipid storage droplet 2* (*lsd2*) have reduced triglyceride stores and a lowered homeostatic sleep response (Thimgan *et al*. 2010). Together, these findings suggest triglycerides, and perhaps energy stores more generally, are required for normal sleep in flies.

The role of adipose tissue in sleep regulation appears to be conserved across phyla. In mammals, leptin is secreted from adipocytes in response to nutritional state and acts on hypothalamic circuits in the brain to decrease feeding as well as increase energy expenditure (Adamantidis and de Lecea 2008; Dardeno *et al*. 2010). In addition, sleep is disrupted in leptin deficient mice, suggesting a role for this adipose-derived hormone in regulating sleep (Laposky 2005). Supporting these findings, our RNAi screen found reduced sleep in flies with fat body knockdown of *upd2*, a proposed *Drosophila* ortholog of mammalian leptin (Rajan and Perrimon 2012). The identification of *Ade2* as well as a number of additional candidate sleep regulators suggest a central role for the fat body in sleep regulation.

*Ade2* encodes a phosphoribosylformylglycinamidine synthase that plays a critical role in purine synthesis in nearly all living organisms (Chaudhary *et al*. 2004). Mutation of *Ade2*, or other components of the *de novo* purine biosynthesis pathways have been implicated in the arrest of cell growth, and reduced fertility and life span, suggesting broad biological functions of this gene (Tiong and Nash 1990; Malmanche and Clark 2004). For example, flies heterozygous for *Ade2* mutations develop necrosis as pupae, suggesting haploinsufficiency may result in physiological abnormalities (Holland *et al*. 2011). The possibility that targeted disruption of *Ade2* in the fat body disrupts development of this organ, or its function in energy storage, is supported by our findings that whole-body triglycerides and/or free glucose levels are reduced in *Ade2* deficient flies. In addition, it is possible that reduced levels of purines themselves contribute to the sleep phenotype. In both flies and mammals, adenosine promotes sleep, and the accumulation of adenosine during periods of wakefulness is associated with increased sleep drive (Jones 2009; Nall *et al*. 2016). Therefore, it is possible that a reduction in adenosine or other purinerginic signaling contributes to the sleep loss phenotype.

Our findings suggest that overexpression of *Ade2* in an otherwise wild type fly does not promote sleep, suggesting *Ade2* is essential for normal sleep, rather than different levels of this gene regulating amounts of sleep. In addition, the partial restoration of sleep to both *Ade2* alleles with fat body specific rescue, suggests *Ade2* may function in the brain or other tissue to regulate sleep. *Ade2* is ubiquitously expressed and it is possible that the haploinsufficiency may be caused by dysregulated purinergic signaling or developmental abnormalities in additional brain regions. In flies, many neural circuits have been found to regulate sleep including the central complex, circadian neurons, and gustatory neurons (Donlea *et al*. 2011; Linford *et al*. 2012; Liu *et al*. 2012; Guo *et al*. 2016). In addition, more recent work has revealed a critical role for non-neuronal tissue, such as glia, in sleep regulation (Chen *et al*. 2015; Farca Luna *et al*. 2017). Targeted disruption of *Ade2* in additional cell types may help reveal novel insights into *Ade2* function.

In flies, starvation results in increased waking activity in addition to reduced sleep duration. We find that waking activity is significantly increased, or trends towards increase, in *Ade2*-deficient flies, indicating that the mutant phenotype recapitulates the hyperactivity induced by starvation. Further, our analysis suggests that *Ade2* is required for normal sleep under baseline conditions, while it is dispensable for sleep homeostasis. Mechanically sleep depriving flies throughout the night results in a rebound the following day, a response that is unaffected in *Ade2*-deficient flies. These data support the notion that independent genetic mechanisms underlie regulation of sleep in undisturbed conditions and during sleep rebound. Along these lines, distinct neural circuits have been identified regulating baseline sleep and recovery sleep in *Drosophila* (Seidner *et al*. 2015a; Liu *et al*. 2016b). Similarly, arousal threshold is unaffected in *Ade2* mutant flies suggesting the quality of the sleep is not dysregulated, but rather the observed phenotype is specific to sleep. Together, these findings suggest *Ade2* regulates baseline sleep duration, but may be dispensable for regulating sleep depth and homeostasis.

A growing body of evidence suggests that the levels of energy storage molecules are critical for normal sleep regulation. Starvation potently reduces whole-body triglycerides and glycogen in *Drosophila* and is associated with reduced sleep (Schwasinger-Schmidt *et al*. 2012). In addition, sleep duration and triglycerides are enhanced in flies selected for starvation resistance (Masek *et al*. 2014). In flies, both sleep and triglycerides levels are sexually dimorphic, with females possessing higher levels of triglycerides and sleeping less during the day (JRD personal observation and (Elwyn Isaac *et al*. 2010)). Feminization of the male fat body through misexpression of the sex differentiation gene *transformer* reduces daytime sleep in males, though the level of triglycerides was not explored (Khericha *et al*. 2016). However, decreasing the expression of the *transformer2* gene in the *Drosophila* fat body results in increased triglyceride storage (Mikoluk *et al*. 2018), suggesting that altering the sex determination pathway in the fly fat body may control both sleep and nutrient storage. Therefore, it is possible that loss of *Ade2* in the fat body induces a chronic starvation-like state, where animals suppress their sleep. Alternatively, the fat body may regulate sleep through a mechanism that is independent of signaling energy stores.

Taken together, the confirmed role of *Ade2* and the identification of additional candidate genes that function within the fat body to either promote or inhibit sleep, supports a central role for the fat body in sleep regulation. The fat body may regulate sleep directly through controlling circulating nutrients that are sensed by the brain or by hormonal communication. In both flies and mammals, sleep modulating neurons directly sense glucose, raising the possibility that circulating glucose levels regulate sleep (Varin *et al*. 2015; Manière *et al*. 2016). The identification of genes that function within the fat body, combined with genetic technology for *in vivo* imaging of sleep circuits, will allow for investigation of how peripheral gene regulation modulates sleep circuits within the brain. The identification of *Ade2*, and other candidate regulators of sleep, provide a platform for investigating the role of periphery-brain communication in sleep regulation.

## Methods

### Fly Stocks

Flies were grown and maintained on standard food (Bloomington Recipe, Genesee Scientific). Flies were kept in incubators (Powers Scientific; Dros52) at 25°C on a 12:12 LD cycle with humidity set to 55-65%. The background control line used in this study is *w^1118^* fly strain, and all experimental flies were outcrossed 6-8 generations into this background, unless already in this background. The following fly strains were ordered from Bloomington Stock Center, w^1118^(5905; Levis *et al*. 1985) and CG-GAL4 (7011; Asha *et al*. 2003). *Ade2^3-20^*, *Ade2^1-6^*, and UAS-*Ade2* were obtained from D. Clark and have been previously characterized (Holland *et al*. 2011). *Drosophila* lines used in the RNAi screen originate from the TRiP collection (Brand and Perrimon 1993; Ni *et al*. 2009) and are described in (Table 2).

### Sleep analysis

The *Drosophila* Activity Monitor System (DAMS) detects activity by monitoring infrared beam crossings for each animal (Pfeiffenberger *et al*. 2010a). These data were used to calculate sleep information by extracting immobility bouts of 5 minutes using the *Drosophila* Counting Macro (Pfeiffenberger *et al*. 2010b; Garbe *et al*. 2015b). For experiments examining the effects of starvation on sleep, flies were kept on 12:12 LD cycle. 5-7 day old female flies were briefly anesthetized with CO_2_ and placed into plastic tubes containing standard food. All flies were given 24 hours to recover after being anesthetized. Activity was recorded for 24 hours on food (ZT0-ZT24).

### Protein, glucose, glycogen and triglyceride measurements

Assays for quantifying triglyceride, glycogen and protein content of flies were performed as previously described (Mikoluk *et al*. 2018). Two bodies from female flies aged 3-5 days were homogenized in buffer containing 50 mM Tris-HCl, pH 7.4, 140mM NaCl, 0.1% Triton-X, 1X protease inhibitor cocktail (Roche). Triglyceride concentration was measured using the Infinity Triglyceride Reagent (ThermoFisher), and protein concentrations were measuring using a BCA Protein Assay Kit (Pierce Scientific). Total glucose levels were determined using the Glucose Oxidase Reagent (Pointe Scientific) in samples previously treated with 8mg/mL amyloglucosidase (Sigma) in 0.2M Sodium Citrate buffer, pH 5.0. Free glucose was measured in samples not treated with amyloglucosidase and then glycogen concentrations were determined by subtracting the free glucose from total glucose concentration. Free glucose, glycogen and triglyceride concentrations were standardized to the total protein content of each sample.

### Sleep deprivation

Five-seven day old fruit flies were loaded into the DAM System and allowed to acclimate for 24 hours. Following acclimation, day sleep (ZT0-ZT12) was measured in undisturbed flies. Flies were then sleep deprived by mechanical stimulation every 2-3 minutes for 12 hours throughout the night time (ZT12-24). The mechanical stimulus was applied using a vortexer (Fisher Scientific, MultiTube Vortexer) and a repeat cycle relay switch (Macromatic, TR63122). Sleep rebound was measured the following day from ZT0-ZT12.

### Arousal Threshold

Arousal threshold was measured using the Drosophila Arousal Tracking system (DART), as previously described (Faville et al. 2015). In brief, individual female flies were loaded in plastic tubes (Trikinectics, Waltham, MA) and placed on plastic trays containing vibrating motors. Arousal threshold was tested with sequentially increasing vibration intensities, from 0 to 1.2 g, in 0.3 g (200 ms) increments, with an inter-stimulus delay of 15 s, once per hour over 24 hours starting at ZT0. Flies were recorded continuously using a USB-webcam (Logitech) at 1 frame per second. The vibrational stimulus and video tracking parameters, and data analysis were performed using the DART interface developed in Matlab (MathWorks, Natick, MA).

### Statistical Analysis

The experimental data are presented as means ± s.e.m. Unless otherwise noted a one-way (ANOVA) followed by Tukey’s post-hoc test was used for comparisons between two or more genotypes and one treatment. Unpaired t-test was used for comparisons between two genotypes. For arousal threshold experiment, the non-parametric Mann Whitney U test was used to compare two genotypes. For two or more genotypes, a Kruskal-Wallis test followed by Dunn’s post hoc. Test. All statistical analyses were performed using InStat software (GraphPad Software 6.0) with a 95% confidence limit (*p* < 0.05).

## Acknowledgements

We would also like to thank Burczyk/Faville/Kottler LTD for the DART system. Carter Burns (FAU) provided technical assistance for *Drosophila*. The authors are grateful to Dr. Denise Clark (University of New Brunswick) for generously providing *Ade2* mutant *Drosophila*. This grant was support by NIH award 1R01 NS085252 to ACK.

---

**Fig S1.**
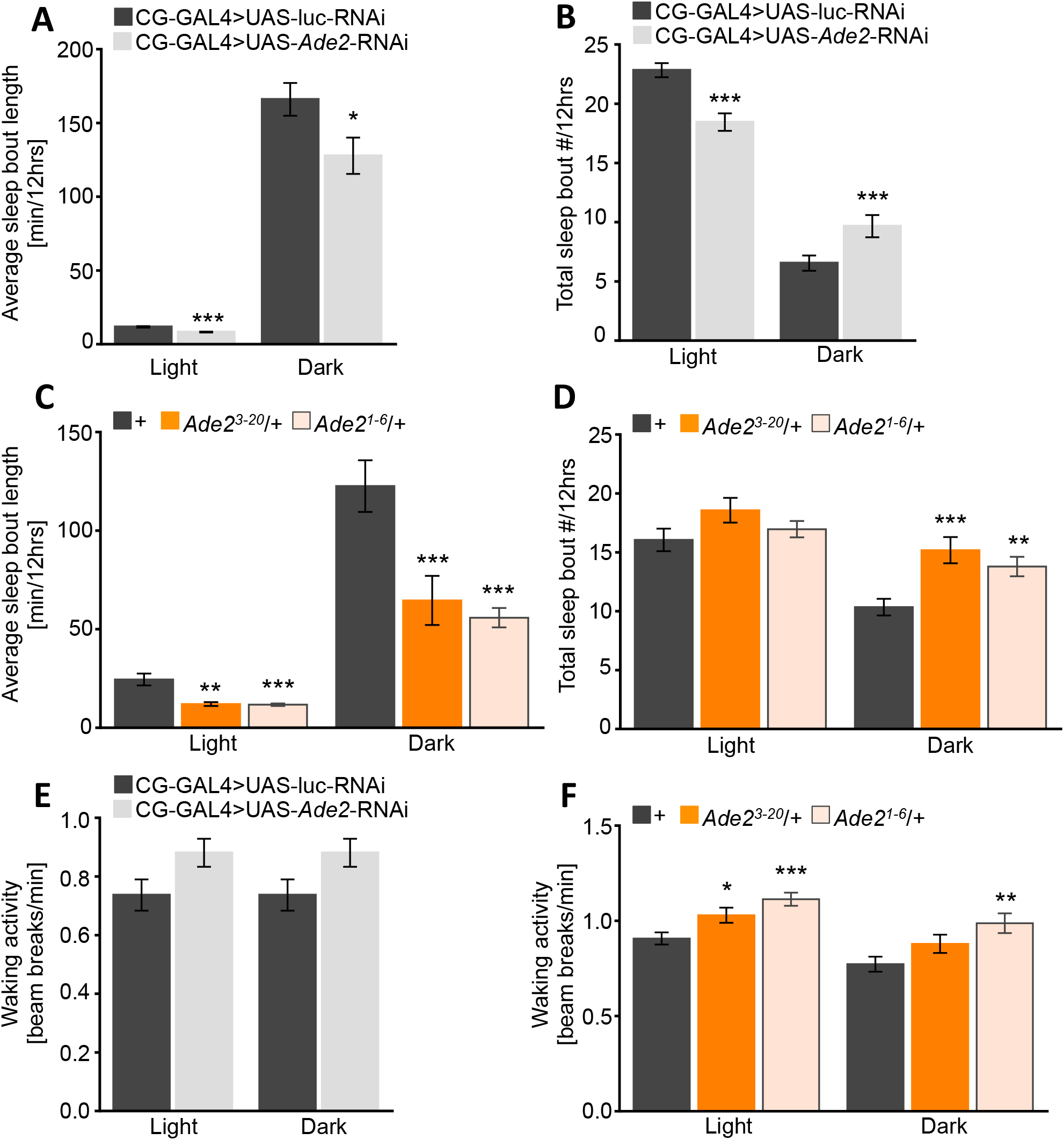
*Ade2* function in the fat body to promote sleep. (**A**) Average bout length is significantly reduced duing dayttime (light; *p*<0.0001, t=5.27) and nighttime (dark, *p*=0.022, t=2.30) in *Ade2-RNAi* (n=87) flies (grey) compared to control (CG-GAL4>UAS-luc-RNAi; black; n=115). Unpaired t-test. (**B**) Knock down of *Ade2* in the fat body (grey) reduced total sleep bout during dayttime (*p*<0.001, t=4.67) compared to control (black), while there is a significant increase during nighttime (*p*=0.005, t=2.83). Unpaired t-test. (**C**) *Ade2^3-20^*/+ (orange; n=61) and *Ade2^1-6^*/+ mutants (pale orange; n=85) reduced average bout length during daytime (*p*<0.0001 for all groups) and nighttime (*p*=0.0019 and *p*<0.0001) compared to *w^1118^* control (black; n=111). One-way ANOVA, Light, F(2, 265)=11.70; Dark, F(2, 254)=11.38. (**D**) Total sleep bout during nighttime is significantly increased in *Ade2^3-20^*/+ (orange; *p*=0.0004) and *Ade2^1-6^*/+ mutants (pale orange; *p*=0.0074) compared to *w^1118^* control (black). One-way ANOVA, F(2, 265)=8.79. (**E**) Waking activity does not differ between control flies (CG-GAL4>UAS-luc-RNAi; black) and *Ade2* knock down in the fat body (grey) during daytime (*p*=0.49, t=0.68) and nighttime (*p*=0.05, t=1.94). Unpaired t-test. (**F**) *Ade2^1-6^*/+ mutants have increased waking activity compared to control flies (*w^1118^*) during daytime (*p*<0.0001) and nighttime (*p*=0.0018), while *Ade2^3-20^*/+ mutants have increased waking activity only during daytime (*p*=0.047). One-way ANOVA; Light, F(2, 265)=9.92; Dark, F(2,265)=6.03. All columns are mean ± SEM; \**p*<0.05; \*\**p*<0.01; \*\*\**p*<0.001.

**Fig S2.**
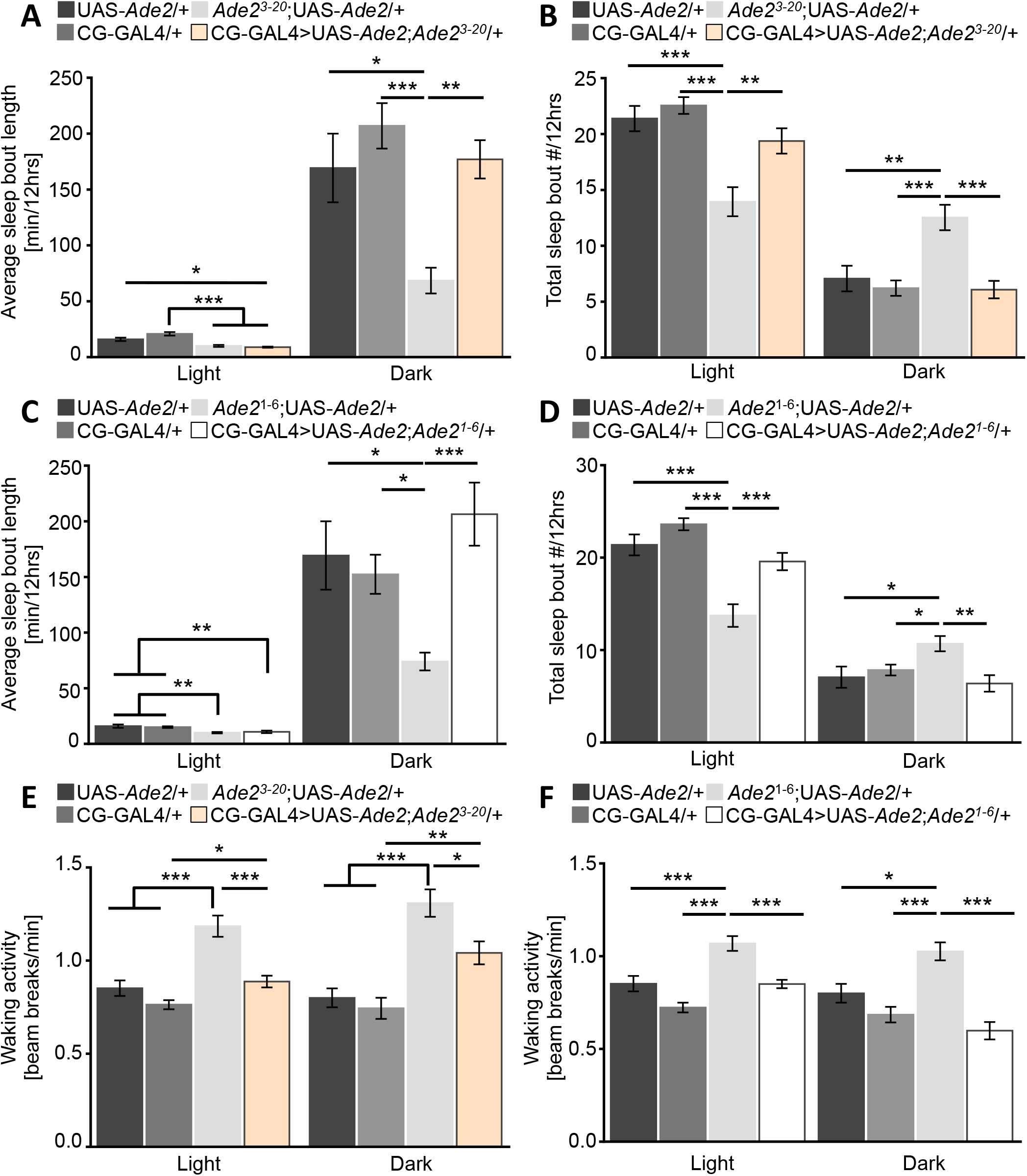
*Ade2* expression in the fat body partially rescues sleep loss. (**A**) Fat body rescue of *Ade2^3-20^* (CG-GAL4>*Ade2^3-20^*;UAS-*Ade2*/+;pale orange;n=50) restores increase in average bout length during the nighttime (dark) compared to *Ade2^3-20^* mutant controls (*Ade2^3-20^*;UAS-*Ade2*/+;grey;n=34, *p*=0.008), but not during dayttime. Daytime average sleep bout length is significantly different between rescue and CG-GAL4/+ (dark grey; n=83, *p*<0.0001) or UAS-*Ade2*/+ (black; n=31, *p*=0.017) controls. Night average sleep bout length is reduced in *Ade2^3-20^*;UAS-*Ade2*/+ compared to control flies CG-GAL4/+ (*p*<0.0001) or UAS-*Ade2*/+ (*p*= 0.036). One-way ANOVA, Light, F(3, 194)=17.76; Dark, F(3,184)=6.865. (**B**) *Ade2^3-20^* rescues total sleep bout during the light (*p*=0.004) and dark (p<0.0001) compared to *Ade2^3-20^*;UAS-*Ade2*/+ mutants. Daytime total sleep bout differ significantly between *Ade2^3-20^*;UAS-*Ade2*/+ mutants and control UAS-*Ade2*/+ and CG-GAL4/+ during light (*p*<0.0001) and dark (*p*<0.0001). One-way ANOVA, Light, F(3, 194)=12.08; Dark, F(3,195)=9.68. (**C**) During nighttime (dark), *Ade2^1-6^* rescue (white, n=43) restores increase in average sleep bout length compared to *Ade2^1-6^* mutant controls (*Ade2^1-6^*;UAS-*Ade2*/+;grey; n=39), but not during daytime. Daytime sleep bout length is significantly different between UAS-*Ade2*/+ (black, n=31, *p*=0.0066) and CG-GAL4/+ (dark grey, n=78, *p*=0.0040) and *Ade2^3-20^* rescue. One-way ANOVA, Light, F(3, 187)=8.98; Dark, F(3,188)=5.412. (**D**) Total sleep bout is restored in *Ade2^1-6^* rescue flies during light (*p*=0.0003) and dark (*p*=0.003) compared to *Ade2^1-6^*;UAS-*Ade2*/+ mutant controls. Daytime total sleep bout is significantly reduced between UAS-*Ade2*/+ and CG-GAL4/+ controls (*p*<0.0001) compared to *Ade2^1-6^* mutant controls, while nighttime sleep bout is increased in *Ade2^1-6^* mutant compared to controls (*p*=0.95). One-way ANOVA, Light, F(3, 187)=21.41; Dark, F(3,184)=4.50. (**E**) Fat body rescue of *Ade2^3-20^* restores waking activity during daytime(*p*<0.0001) and nightime (*p*=0.044) compared to *Ade2^3-20^*;UAS-*Ade2*/+ mutant controls. Waking activity is significantly increased during light (*p*<0.0001) and dark (*p*<0.0001) in *Ade2^3-20^* mutant compared to UAS-*Ade2*/+ and CG-GAL4/+ controls. One-way ANOVA, Light, F(3, 194)=23.79; Dark, F(3,196)=14.34. (**F**) Waking activity is rescued in CG-GAL4>*Ade2^1-6^*;UAS-*Ade2*/+ during daytime (*p*<0.0001) and nightime (*p*<0.0001) compared to *Ade2^1-6^*;UAS-*Ade2*/+ mutant controls. *Ade2^1-6^* mutant control have increased waking activity during light (*p*<0.0001) and dark (*p*=0.02) compared to control flies. One-way ANOVA, Light, F(3, 187)=21.47; Dark, F(3,188)=13.23. All columns are mean ± SEM; \**p*<0.05; \*\**p*<0.01; **-\**p*<0.001.

